# Could microbes be the architects of improved soil structure under *Miscanthus* × *giganteus*?

**DOI:** 10.64898/2026.08.06.743358

**Authors:** Phillip de Lorimier, Jessica T. Nelson, Bolívar Aponte Rolón, Jared Flater, Lorien Radmer, Marshall D. McDaniel, Adina Howe

## Abstract

The perennial grass *Miscanthus* × *giganteus* (miscanthus) offers a sustainable alternative to traditional biomass feedstocks while improving key soil health parameters, including aggregation. Aggregate stability results from dynamic soil-plant-microbe interactions, yet the relative importance of each factor remains an active research question. Building on previous observations that miscanthus alters soil structure to improve water-holding capacity and aggregate stability, we characterized the communities of soil bacteria and arbuscular mycorrhizal fungi (AMF) across three sites in Iowa, USA, comparing miscanthus to annual maize (*Zea mays* L.) and non-cropped perennial turfgrass (*Poa* spp.). We examined whether microbiomes co-varied with soil aggregation and, if so, whether plant cover identity or life history categorization better explained the observed patterns. Bacterial and AMF communities varied across sites and plant types, with signals that life history and plant cover identity both mattered. Aggregate stability aligned with a perennial-annual divergence in microbial beta diversity, while finer-scale differences in community composition and network structure were plant-specific. Soils under perennial plants were enriched in microbial groups positively correlated with aggregate stability; we identified 61 bacterial and 8 AMF “architect” taxa for future study. Within- and cross-kingdom co-occurrence network analysis revealed greater complexity under perennial plants: 1.9-fold more network links in miscanthus bacteria-bacteria networks than in maize, and 1.7-fold more in turfgrass AMF-AMF networks. Miscanthus fundamentally shapes microbial interactions, particularly among bacteria, relating to improved soil physical structure. Understanding these soil-plant-microbe feedbacks advances the development of biomass feedstocks with a portfolio of soil health benefits for next-generation biofuels and bioproducts.

**IMPORTANCE:** Perennial bioenergy crops can provide the raw material for biofuels and bioproducts while simultaneously improving soil health. *Miscanthus* × *giganteus* (miscanthus) efficiently stabilizes soil aggregates, potentially leading to higher water retention and erosion resistance. Understanding the microbial contributions to these outcomes is key to building resilient, sustainable bioenergy systems. This study highlights the connections between communities of soil microbes—bacteria and arbuscular mycorrhizal fungi—across three sites and three plant covers, including miscanthus, maize, and turfgrass. We identify a guild of potential “microbial architects” linked to soil aggregation and show more interconnected microbial networks under the perennial plant covers compared to annual maize. These insights shed light on the interactions between soil biological communities and soil physical and chemical properties. More broadly, the results may inform efforts to harness plant-associated microbiomes for sustainable biomass production.

## 1. INTRODUCTION

Soil aggregation emerges from soil-plant-microbe interactions, and crop attributes govern these feedbacks in agricultural systems (1, 2). First, the crop roots themselves act as a nexus of soil aggregation; their morphology and longevity regulate microbial niches (3, 4). Rhizodeposition fuels microbial production of extracellular polymeric substances (EPS) that can bind soil particles into larger aggregates, improving pore connectivity (5, 6). Compared to conventional annual crops, where this process resets each season, perennial grasses sustain year-round root stocks and prolong rhizodeposition. This, in turn, enriches microbial activity and potentially accelerates aggregate formation and stabilization (7, 8). These dual benefits—restoring soil structure while producing economically viable yields—have spurred interest in perennial biomass crops and underscore the need to elucidate the belowground interactions that sustain them.

The perennial grass *Miscanthus* × *giganteus* (miscanthus) offers distinct advantages as a biomass feedstock. Miscanthus produces high biomass yields and is tolerant of flooded, saline, and low-fertility soils (9, 10). Compared to conventional annual crops such as maize, miscanthus requires fewer inputs, like nitrogen fertilizer (11–13). Another advantage (and unique feature) of miscanthus is that it partitions substantial amounts of unharvested plant material to soil organic matter. This occurs belowground with its rhizome-root complex and aboveground with a distinctive litter layer that can accumulate to 2-4 cm in less than 10 years (14, 15). These two pathways likely contribute to observations of soil carbon (C) sequestration on the order of 0.4-3.2 Mg C ha^−1^ y^−1^ (16–19).

Inputs of root and shoot C are one of many unique characteristics of miscanthus that likely structure a belowground microbial community characterized by high diversity and biomass (20–22). Previous studies have characterized miscanthus microbiomes as temporally stable, though community composition is still shaped by stand age and nutrient availability (23). Miscanthus grown in nutrient-poor soil tends to enrich arbuscular mycorrhizal fungi (AMF) and nitrogen-fixing bacteria that support its nutrient acquisition (24). Additionally, miscanthus rhizodeposition results in greater hydrolytic enzyme activity and C use efficiency compared to annual crops (25, 26). Most previous efforts have focused on the soil bacteria under miscanthus, whereas AMF-miscanthus relationships are relatively under-characterized despite their potential role in nutrient acquisition, as well as in soil aggregate formation and stabilization.

Cultivating miscanthus, even for just a few years, can alter soil structure in ways that enhance both water storage and resistance to erosion (27–29). For example, miscanthus grown for seven years can substantially increase maximum water holding capacity and stabilize soil aggregates (29, 30). Gains in soil aggregation tend to generate other soil health benefits, including increased water retention (31, 32), reduced erosion (33), greater carbon sequestration (34, 35), and greater plant-nutrient supplying potential (36). A basic research question, however, remains: how does miscanthus improve soil structure so quickly compared to other perennial cropping systems?

Nelson et al. (2025) developed a framework of *direct* and *indirect* mechanisms that may contribute to improved soil structure and increased water holding capacity under miscanthus (Figure S2). Briefly, the direct pathways encompass root morphology, rhizodeposition, and other biophysical interactions between miscanthus and the soil matrix (excluding microbes or soil fauna). Indirect pathways include the secondary effects of miscanthus characteristics that shape soil microbial communities, whose activity improves soil structure through EPS production (37), enhanced hyphal production (38), or greater abundance of meso-/macro-fauna (39). We have recent evidence showing that miscanthus soils contained lower ratios of microbe-derived to plant-derived carbohydrates across three sites (30). This enrichment in plant-derived chemicals suggests a greater influence of the *direct* pathway; however, the *indirect* role of soil microbes under miscanthus also merits attention.

Considering this framework and the importance of microbes as soil architects that restructure the soil environment around them, it remains unclear whether improvements in soil structure are a consequence of miscanthus’ perennial life cycle or whether it supports a distinct soil microbiome compared to other perennial or annual plant covers. In particular, the extent to which bacterial and AMF communities differ between miscanthus, annual maize, and other perennials, and whether such differences correspond to variation in soil aggregation, remains largely uncharacterized. Therefore, our first objective was to characterize soil bacterial and AMF communities and quantify their variation across sites and plant covers. We hypothesized that the differences in soil structural properties between plant covers would correspond to variation in microbial community composition. Our second objective was to determine whether the relationships between soil aggregate stability and bacterial and AMF communities were consistent across sites and plant covers. Specifically, we identified taxa associated with aggregate stability and used network analyses to evaluate patterns of microbial co-occurrence and potential interactions, asking whether miscanthus supports distinct microbial assemblages and microbe-microbe associations relative to annual and perennial alternatives.

## 2. MATERIALS AND METHODS

### 2.1 Site Description and Experimental Design

Soil samples were collected in 2022 from the Long-term Assessment of Miscanthus Productivity and Sustainability (LAMPS) experiment across three sites in Iowa, USA: Northwest, Central, and Southeast (40). Although coming from two landform regions of Iowa, all soils developed under glacial till or a mix of glacial till and loess cap (41). Soils at the Southeast site are predominantly silty loams, whereas the Central and Northwest sites have higher clay content and are classified as clay loams (Table 1).

**Table 1.**
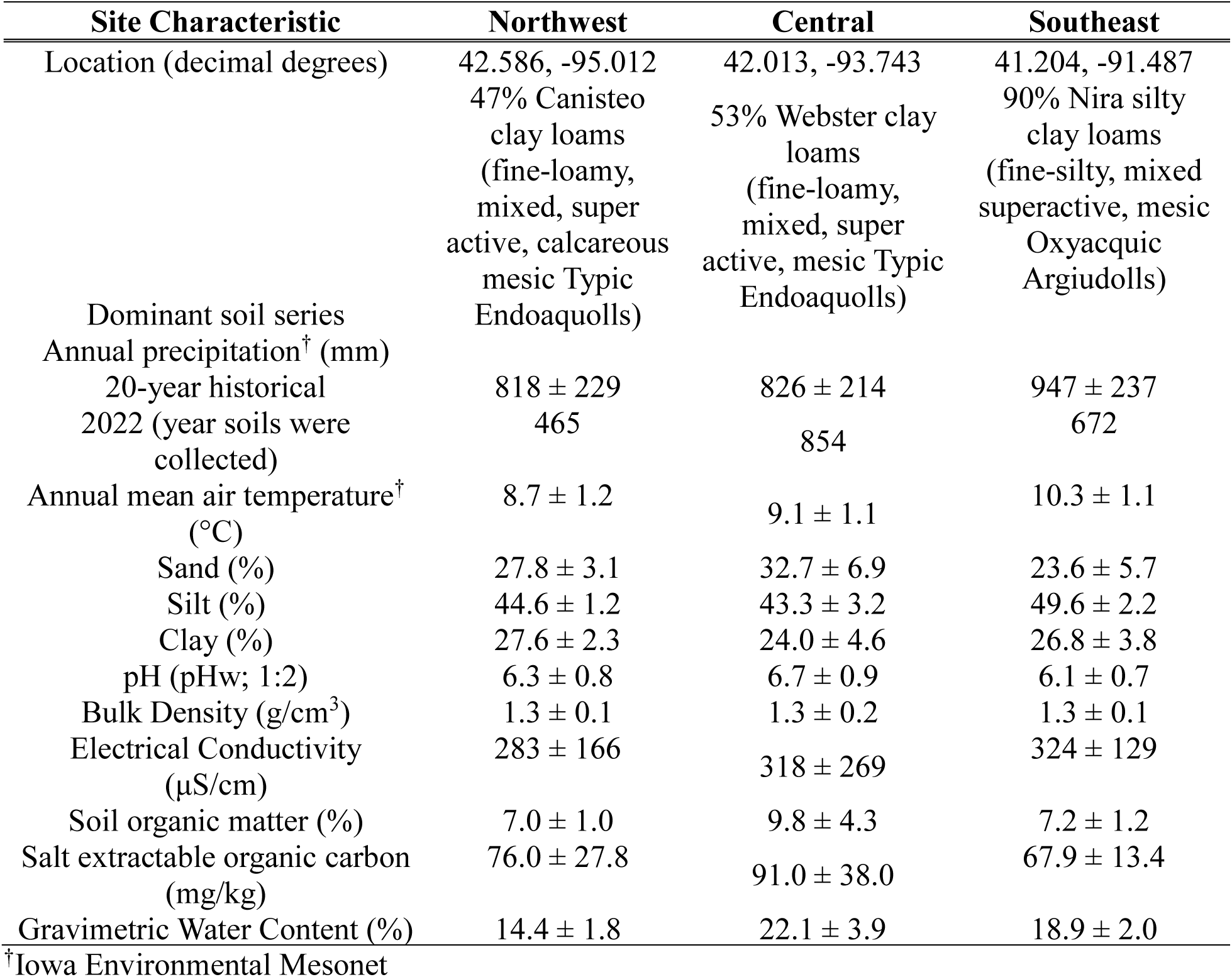
Characteristics for the Northwest, Central and Southeast, Iowa, USA sites in the Long-Term Assessment on Miscanthus Productivity and Sustainability (LAMPS) experiment (n = 12 per site; mean values ± SD).

**Table 2.** Summary of sample counts, total amplicon sequence variants (ASVs) and total unique ASVs observed within each treatment.

| **Marker** | **Treatment** | **Samples** | **Total ASVs** | **Unique ASVs** |
| --- | --- | --- | --- | --- |
| Bacteria | Maize | 12 | 3503 | 250 |
| Bacteria | Miscanthus | 12 | 3939 | 608 |
| Bacteria | Turfgrass | 12 | 3336 | 395 |
| AMF | Maize | 12 | 322 | 192 |
| AMF | Miscanthus | 11 | 200 | 99 |
| AMF | Turfgrass | 10 | 271 | 133 |

All sites had been in a maize-soybean rotation with conventional tillage before the experiment’s initiation in 2015. At that time, miscanthus was planted, and a continuous maize rotation was implemented for the maize treatment. The maize plots were managed with conventional tillage until 2017, when they were converted to no-till, and the miscanthus plots have been managed with no-till since establishment. The Northwest site received annual manure applications containing ∼50% cornstalks, applied at approximately 11,208 kg ha⁻¹ y⁻¹, supplying an estimated 4,537 kg C and 154 kg N ha⁻¹ y⁻¹. After miscanthus was planted in 2015, this practice was replaced by synthetic nitrogen (N) application. The Central and Southeast sites were managed solely with synthetic fertilizers for at least 20-30 years.

The study was a split-plot, randomized block design with four replications per cropping system. Main plots were 24 × 60 m, with split-plots measuring 24 × 12 m. Miscanthus plots were planted in 2015 and fertilized annually at 224 kg N ha^−1^. We sampled only the 2015 miscanthus plantings, which were in their seventh growing season at the time of sampling (October 2022). We sampled from the continuous maize plots, which were fertilized annually at 224 kg N ha^−1^. We also sampled from the bordering grass alleyways, where there was minimal heavy machinery traffic at all three sites, for comparison with perennial, short-statured turfgrasses such as fescue (*Festuca arundinacea*), Kentucky bluegrass (*Poa pratensis*), and foxtail (*Setaria viridis*). Hence, the study involved four replications for three plant types (maize, miscanthus, turfgrass), resulting in twelve experimental units at all three sites (Northwest, Central, Southeast).

### 2.2 Soil Sampling, Ancillary Measurements, and Pre-processing for DNA Extraction

Within each plot, 10 soil cores (1.75 cm diameter, 15 cm depth) were collected in a stratified random pattern to avoid plot boundaries while remaining representative of the plot, and then thoroughly mixed to create a composite sample. The soil was then sieved at field moisture through 2- and 8-mm sieves to retain two size classes: i) < 2 mm to homogenize for fresh soil analyses and ii) 2-8 mm sizes to retain larger aggregates for aggregate stability testing.

Laboratory analyses were performed for a wide range of soil physical and biochemical properties, previously described in (30), to evaluate differences in soil structure, water retention, and biochemical signatures. These measurements include bulk density, root density, soil organic matter, aggregate stability, penetration resistance, gravimetric water content, maximum water holding capacity, microbial biomass carbon, salt-extractable organic carbon, EPS, and soil carbohydrates. A summary of these measurements is provided in the Supplementary Information (Table S1).

### 2.3 DNA extraction and amplicon sequencing

For DNA extraction, a 0.25 g subsample was taken from the portion of each sample sieved to < 2 mm. DNA extraction was performed using the MagAttract PowerSoil DNA EP kit (Qiagen, Germantown, MD, USA) according to the kit’s standard protocol, with an Eppendorf epMotion 5075 liquid-handling instrument (Eppendorf, Enfield, CT, USA). DNA samples with a concentration above 10 ng μl^−1^ were diluted to 10 ng μl^−1^ before sequencing. Samples with concentrations lower than 10 ng μl^−1^ were submitted directly for amplicon sequencing.

The V4 region of the bacterial 16S ribosomal RNA (rRNA) gene was amplified for the community analysis of bacteria. Amplification was performed using 10 µM each of 16S rRNA v4 region primers (42). The forward primer (515F) sequence was GTGYCAGCMGCCGCGGTAA, and the reverse primer (806R) sequence was GGACTACNVGGGTWTCTAAT. The target amplicon size was 390 bp. The PCR amplification parameters were as follows for a 384-well plate: 94°C for 3 min; then 94°C for 60 s, 50°C for 60 s, and 72°C for 105 s, repeated for 35 cycles, with a final extension at 72°C for 10 min.

A region of the 18S ribosomal RNA (rRNA) gene was amplified for the AMF community analysis. Amplification was performed using 10 µM each of 18S rRNA primers (43). The forward primer (NS31) sequence was TTGGAGGGCAAGTCTGGTGCC, and the reverse primer (AML2) sequence was GAACCCAAACACTTTGGTTTCC.

Sequencing of bacterial and AMF amplicons was performed on an Illumina MiSeq using the MiSeq Reagent Kit V2 (Illumina, San Diego, CA, USA) at Argonne National Laboratory (Argonne, IL, USA), and the libraries consisted of 150-bp paired-end reads. Sequencing data are deposited in the NCBI Short Read Archive as project PRJNA1366626.

### 2.4 Bioinformatic and Statistical Analyses

The sequencing data were analyzed using the DADA2 package (version 1.14.0) to determine the abundance of amplicon sequence variants (ASVs) (44). For bacterial amplicons, the truncated sequence read length was set to 145 bp to remove low-quality tails based on inspection of quality control profiles. The filtering parameters were: truncLen = c(145,145), truncQ = 2, rm.phix = TRUE, and compress = TRUE. The taxonomic identity of each observed ASV was determined using sequence similarity to representatives in the SILVA rRNA database version 138.1. ASVs observed in < 2 samples were removed. For AMF amplicons, sequences were not truncated to a fixed length due to the greater variability in the 18S rRNA gene region. Primers were removed using *cutadapt()* version 5.0, and the taxonomic identity of each observed ASV was determined using sequence similarity to representatives in the MaarjAM database (45).

All statistical analysis was performed in R version 4.4.1 (46). Permutational multivariate analysis of variance (PERMANOVA) was performed with the *adonis2()* function from the vegan package, version 2.6-10 (47), based on Bray-Curtis dissimilarity distances between samples, with p-values for the test statistic (pseudo-F) obtained via 999 permutations (Table 3). Principal coordinates analysis was performed to visualize sample dissimilarity using the Bray-Curtis dissimilarity matrix. To assess the influence of soil physicochemical variation on the microbial community composition, we conducted a distance-based redundancy analysis (dbRDA) using the *dbrda()* function from the vegan package. The principal coordinates from the previous analysis were fitted to a multiple linear regression using bulk density, water-stable aggregate percentage, root density, gravimetric water content, and EPS concentration as explanatory variables. These variables were chosen to test the hypothesis that soil physical and chemical properties are associated with differences in microbial community composition. The resulting fitted values and residuals were processed using separate principal component analyses to produce an ordination that maximized the fit between the constraining environmental variables and the original distance matrix (Table 4).

**Table 3.** Results from PERMANOVA test of bacterial and arbuscular mycorrhizal fungal beta diversity.

| **Marker** | **Factor** | **Degrees of Freedom** | **Sum Of Squares** | **R<sup>2</sup>** | **F Statistic** | **p-value** |
| --- | --- | --- | --- | --- | --- | --- |
| Bacteria | Site | 2 | 1.3963 | 0.1581 | 3.397 | **0.001** |
| Bacteria | Treatment | 2 | 0.9168 | 0.1038 | 2.2305 | **0.005** |
| Bacteria | Site:<br>Treatment | 4 | 0.9676 | 0.1096 | 1.177 | 0.179 |
| Bacteria | Residual | 27 | 5.5489 | 0.6284 |  |  |
| AMF | Site | 2 | 1.3699 | 0.1078 | 2.0477 | **0.001** |
| AMF | Treatment | 2 | 1.4809 | 0.1166 | 2.2136 | **0.001** |
| AMF | Site:<br>Treatment | 4 | 1.8274 | 0.1438 | 1.3658 | **0.003** |
| AMF | Residual | 24 | 8.0279 | 0.6318 |  |  |

**Table 4.** Summary of distance-based redundancy analysis (dbRDA) model summary, overall constrained variance (%) and the F statistics of individual constraining variables (See Supplemental Information for details on measurements).

| **Component (or**<br>**soil**<br>**measurement)** | **Bacteria** |  | **Arbuscular Mycorrhizal Fungi (AMF)** |  |
| --- | --- | --- | --- | --- |
|  | **F Statistic** | **p-value** | **F Statistic** | **p-value** |
| Overall model | 1.6302 | **0.003** | 1.5087 | **0.001** |
| Bulk Density | 2.7540 | **0.001** | 1.7193 | **0.004** |
| Water Stable<br>Aggregate % | 1.4923 | 0.103 | 1.6006 | **0.008** |
| EPS | 0.6219 | 0.955 | 0.8018 | 0.853 |
| Root Density | 1.0328 | 0.344 | 1.2543 | 0.126 |
| GWC | 1.0937 | 0.303 | 1.9744 | **0.002** |
| MWHC | 2.7868 | **0.002** | 1.7016 | **0.005** |

Spearman correlations were estimated between the abundance of taxa and the environmental measurements, and we filtered for taxa where the correlation with key physical properties—water-stable aggregate %, mean weight diameter, or maximum water-holding capacity—was at least moderately positive (Spearman rho > 0.4, *p* < 0.05). We compared the relative abundance of these taxa across sites and treatments. Finally, to assess the effect of perenniality, we computed the log_2_-fold change of each taxon’s abundance from the annual system (maize) to the mean value of the perennial systems (miscanthus and turfgrass) at each site, adding a trivial pseudocount (1 × 10^−6^) to all values to avoid an undefined calculation.

We constructed bacteria-bacteria, AMF-AMF, and bacteria-AMF co-occurrence networks to analyze inter- and intra-kingdom ecological relationships, following the approach of (48). For the intra-kingdom networks, we retained ASVs that were present in at least two samples. Next, we calculated pairwise correlations among all ASVs using the Pearson correlation of their relative abundances across the 12 replicates. Pairs of ASVs with a correlation above 0.4 were considered “linked” in the network. For the inter-kingdom networks, the ASV tables for bacteria and AMF were first combined to represent a single community before the correlation steps were performed. The resulting networks were visualized using Gephi version 0.10.1. To assess the potential ecological role of the aggregate-correlated ASVs identified in the previous analysis, we filtered the network link tables to include only links containing an aggregate-correlated bacterial or AMF ASV, and the total number of links between treatments was compared with the *t.test()* function in R.

Code for all described analyses is available at https://github.com/germs-lab/LAMPS_EPS_manuscript.

## 3. RESULTS

### 3.1 Bacterial and AMF communities, and their relationship to soil properties

Across a total of three sampling sites and three crops, we observed a total of 5,131 bacterial ASVs classified to 29 phyla and 584 AMF ASVs classified to 7 genera within the phylum *Mucoromycota* (Table 2, Figure S3). Microbiome membership within each plant cover consisted of both unique and broadly shared ASVs: 76% of bacterial ASVs and 27% of AMF ASVs were detected under more than one plant cover (Figure S4). Beta diversity was assessed using Bray-Curtis dissimilarity to quantify differences in microbial community composition between plant covers and sites. PERMANOVA analysis revealed that for bacteria, the primary factor affecting this diversity was the site, followed by plant type (Table 3, *p* < 0.001). There was no evidence of an interaction between site and plant type on bacterial beta diversity. For AMF, the difference in beta diversity due to plant type explained slightly more variation than the site, and there was evidence for an interaction between plant type and site (Table 3, *p* < 0.05).

For the bacterial community, the constrained dbRDA ordination revealed that soil physical and chemical properties explained 25.2% of the total variance in community composition (Figure 1a). ANOVA-like permutation tests indicated that bulk density (*p* = 0.008) and maximum water holding capacity (*p* = 0.005) significantly constrained the ordination (Table 4). For AMF, 25.8% of the total variance in community composition was constrained by the measured soil physical and chemical properties (Figure 1b). Permutation tests for the dbRDA indicated that bulk density (*p* = 0.004), water-stable aggregate % (*p* = 0.002), gravimetric water content (*p* = 0.001), and maximum water-holding capacity (*p* = 0.005) significantly constrained this ordination (Table 4). Notably, the dbRDA ordination indicated that perennial systems (miscanthus and turfgrass) exhibited greater similarity to each other in microbial community composition than they did to the annual maize system.

**Figure 1.**
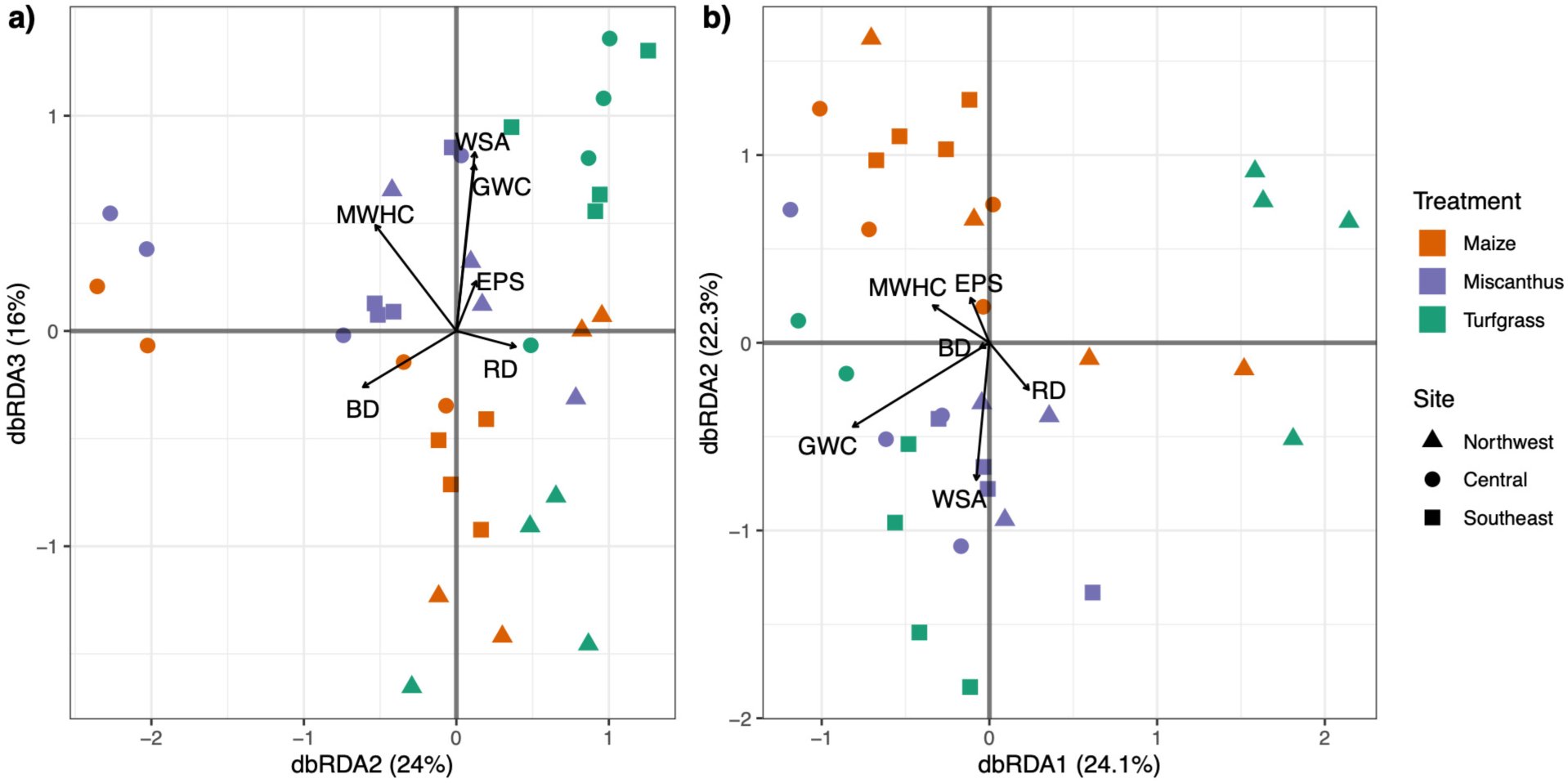
Distance-based redundancy analysis (dbRDA) showing the association of key soil physical and chemical properties with **(a)** bacterial and **(b)** arbuscular mycorrhizal fungal beta diversity. Abbreviations: BD: bulk density, WSA: water-stable aggregates, EPS: extracellular polymeric substances, RD: root density, GWC: gravimetric water content, MWHC: maximum water holding capacity. See Table 4 for summary statistics of the dbRDA.

### 3.2 Characterizing microbial taxa with linkages to soil aggregation

Bacterial and AMF taxa correlated with aggregate stability were enriched in miscanthus and turfgrass soils relative to maize. We identified taxa in our samples that may be linked to soil aggregation and maximum water holding capacity. This resulted in the identification of 61 bacterial ASVs and 8 AMF ASVs, which we refer to as putative “microbial architects” (Table S2). Considering miscanthus and turfgrass together as perennial treatments and comparing them to annual maize on a site-by-site basis, we found that aggregate-correlated bacterial ASVs were substantially more enriched under miscanthus and turfgrass (Figure 2a). Notably, the aggregate-correlated ASVs were overwhelmingly detected in samples spanning multiple sites in both the miscanthus and turfgrass treatments (Figure 2b). The magnitude of enrichment or depletion varied considerably, with an average mean log_2_-fold change of 3.7 (range: −1.8 to 9.7). Aggregate-correlated AMF taxa also tended to be enriched in perennial relative to annual systems, with a mean log_2_-fold change of 3.3 (range: 0 to 7.0) (Figure 3a). Overall, the bacterial taxa correlated with aggregate stability exhibited higher relative abundance under miscanthus and turfgrass than in maize, a pattern observed across the three sites in this study, though most pronounced at the Southeast site (Figure 4a). The relative abundance of aggregate-correlated AMF taxa was highly site-specific; here again, the Southeast site showed the strongest perennial enrichment of ASVs in the perennial plant covers (Figure 4b).

**Figure 2.**
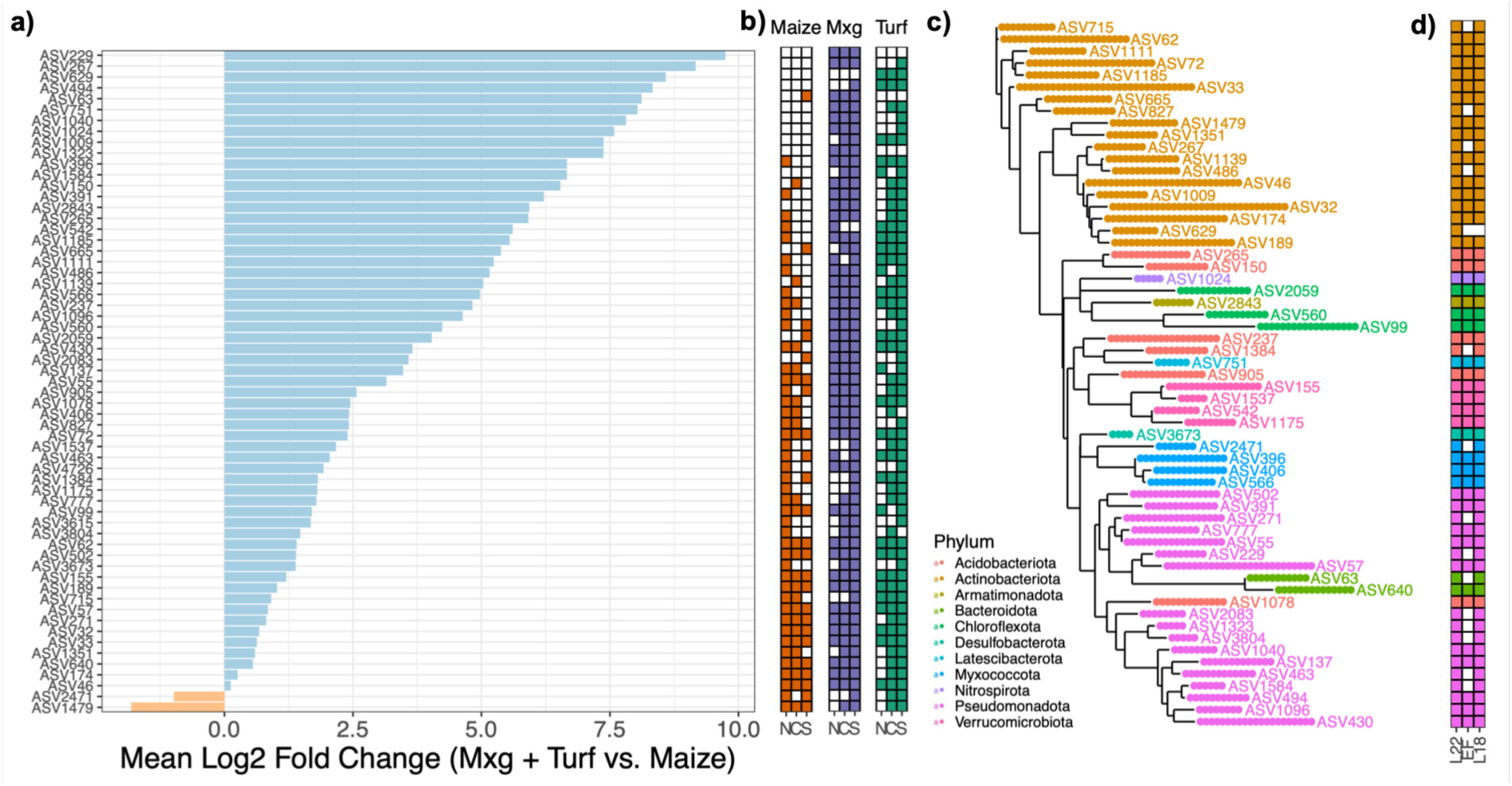
Characterization of aggregate-correlated bacterial ASVs (Spearman *ρ* > 0.4, p < 0.05). **(a)** Mean Log_2_-fold change in the abundance between perennial (miscanthus and turfgrass) and annual (maize) systems. **(b)** Presence (filled square) or absence (empty square) of each ASV within sites (‘N’: Northwest, ‘C’: Central, ‘S’: Southeast) for maize, miscanthus (‘Mxg’), turfgrass (‘Turf’). (**c)** Maximum-likelihood phylogenetic trees based on nucleotide similarity of aggregate correlated ASVs. **(d)** Presence (filled square) or absence (empty square) of each ASV in comparison datasets representing a different location (‘EF’: Energy Farm, Urbana, IL) or at the same site at an earlier timepoint (‘L18’: LAMPS 2018) compared to this present study (‘L22’: LAMPS 2022).

**Figure 3.**
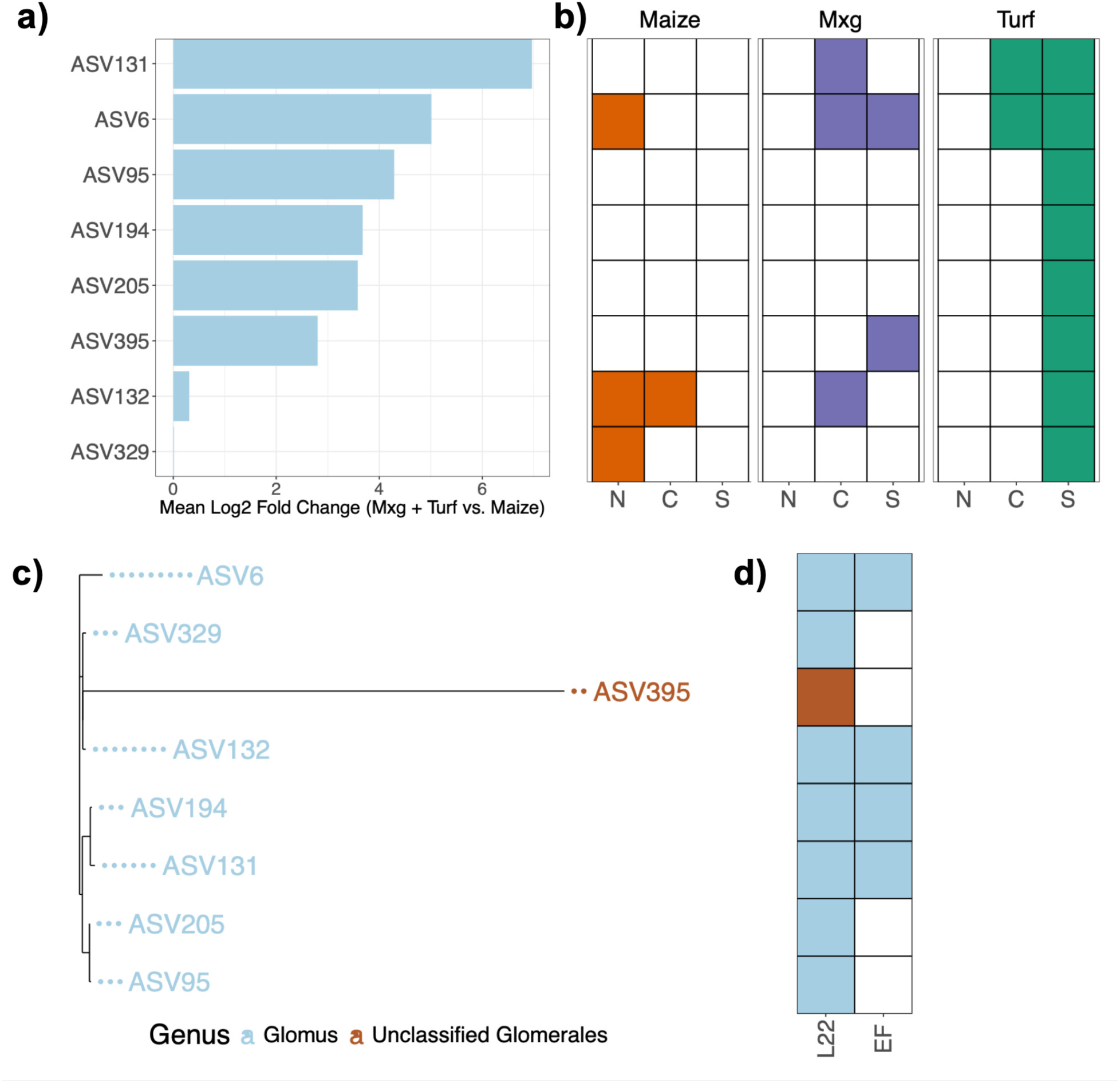
Characterization of aggregate-correlated AMF ASVs (Spearman *ρ* > 0.4, p < 0.05). **(a)** Mean Log_2_-fold change in the abundance between perennial (miscanthus and turfgrass) and annual (maize) systems. **(b)** Presence (filled square) or absence (empty square) of each ASV within sites (‘N’: Northwest, ‘C’: Central, ‘S’: Southeast) for maize, miscanthus (‘Mxg’), turfgrass (‘Turf’). (**c)** Maximum-likelihood phylogenetic trees based on nucleotide similarity of aggregate correlated ASVs. **(d)** Presence (filled square) or absence (empty square) of each ASV in comparison datasets representing a different location (‘EF’: Energy Farm, Urbana, IL) compared to this present study (‘L22’: LAMPS 2022).

**Figure 4.**
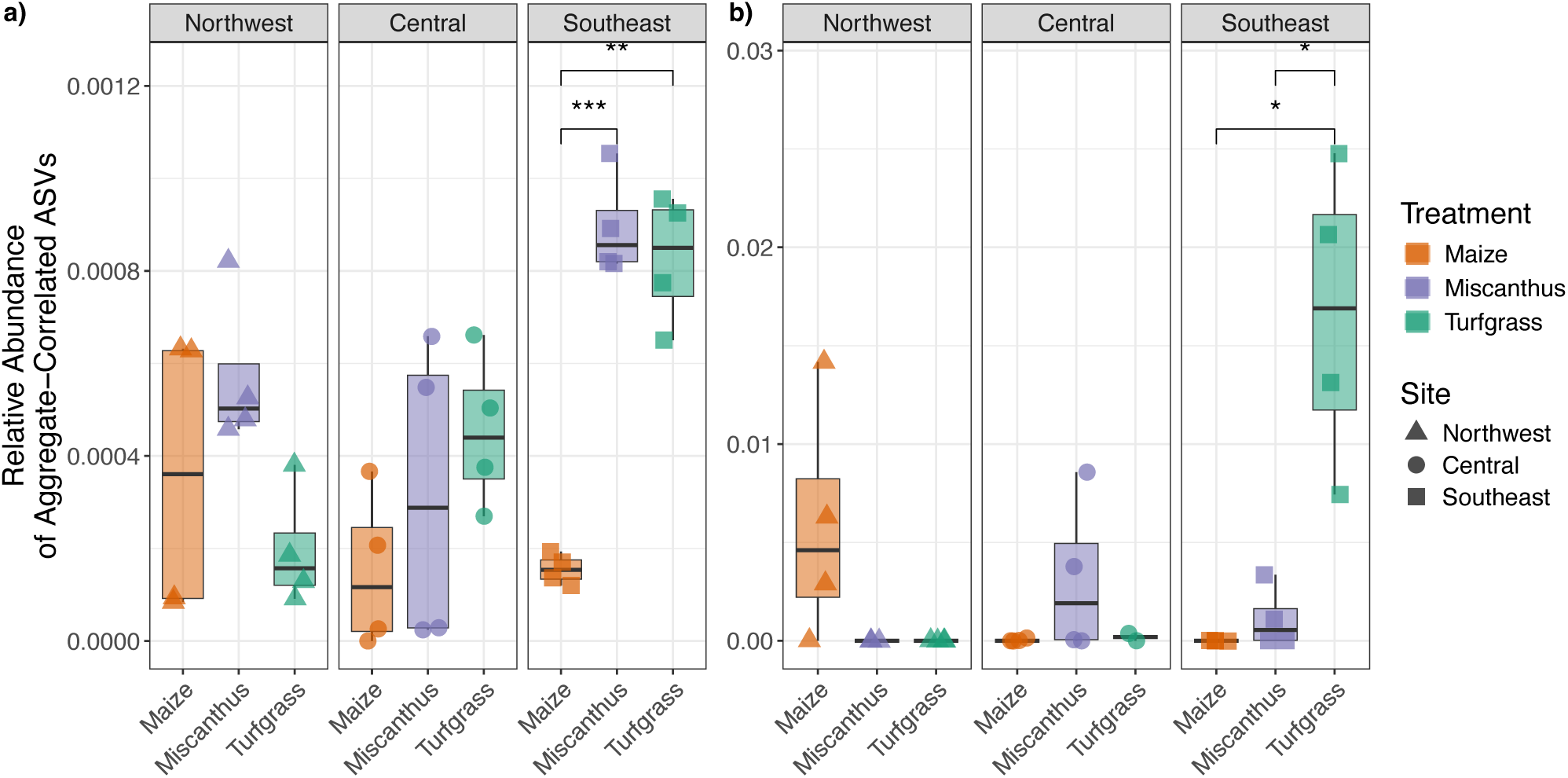
Relative abundance of **(a)** bacterial and **(b)** arbuscular mycorrhizal taxa significantly correlated with water stable aggregates (proportion of > 2mm aggregates remaining intact following wet sieving). Individual points are plot scale. Significant differences within site and among plant cover means are shown: * p ≤ 0.05; ** p ≤ 0.01; *** p ≤ 0.001.

Considering the taxonomy of aggregate-correlated bacterial ASVs, the dominant phyla were *Actinobacteriota* (32%) and *Pseudomonadota* (29%) (Figure 2c). Of the eight AMF ASVs, seven were classified to the genus *Glomus*, and one was an unclassified member of the order *Glomerales* (Figure 3c). Finally, to assess whether the aggregate-correlated ASVs identified in this study were specific to our experimental conditions or represented broader patterns, we evaluated three independent miscanthus microbiome datasets: bacterial and AMF communities from miscanthus soils in Urbana, Illinois collected during the 2023 growing season, and bacterial communities from the Central Iowa site collected in 2018, four years before the sampling for this study. ASVs with >99% sequence similarity were considered a match. Bacterial ASVs were consistently detected across sites and time: 49 out of 61 aggregate-correlated ASVs were detected in the Illinois dataset, and nearly all were detected at the Central Iowa site in 2018 (Figure 2d). AMF ASVs were less consistent across sites: 50% of the aggregate-correlated ASVs were detected in the Illinois dataset (Figure 3d).

### 3.3 Network analysis of aggregate-correlated taxa to explore cross-domain relationships

By compiling aggregate-correlated microbe-to-microbe networks within maize, miscanthus, and turfgrass soils, we found that miscanthus and turfgrass soils were generally characterized by significantly more complex networks than maize soils (Figure 5a-c). Notably, miscanthus bacteria-bacteria networks had 1.9-fold and 1.6-fold more network links than maize and turfgrass, respectively (Figure 5d). Turfgrass had the most AMF-AMF network links, with 5.7-fold and 1.7-fold more than miscanthus and maize, respectively (Figure 5f). Bacteria-AMF interactions were similar across all three plant covers (Figure 5e). Between the treatments, no differences in within-module connectivity (zi) or among-module connectivity (pi) were detected (Table S3).

**Figure 5.**
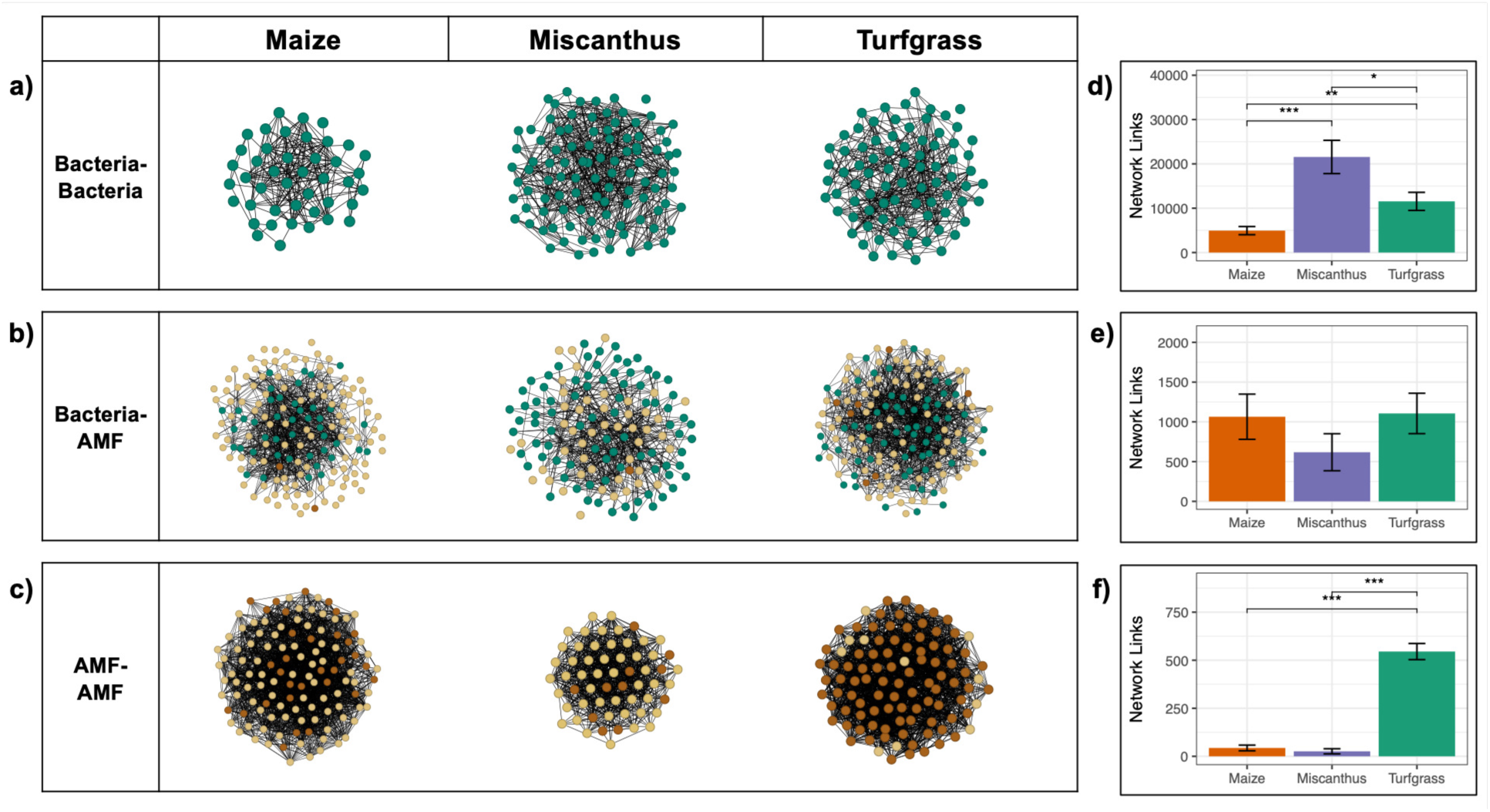
Central log-ratio Pearson cooccurrence network structures for **(a)** bacteria-bacteria, **(b)** bacteria-AMF, and **(c)** AMF-AMF under maize, miscanthus, and turfgrass. Blue points indicate bacterial taxa correlated with water stable aggregate %, dark brown points indicate AMF taxa correlated with water stable aggregate %, and light brown points indicate all other AMF taxa. Bar plots showing mean and standard error (n = 12) of network links for **(d)** bacteria-bacteria, **(e)** bacteria-AMF, and **(f)** AMF-AMF connections. Significant differences among plant cover means are shown: * p ≤ 0.05; ** p ≤ 0.01; *** p ≤ 0.001.

## 4. DISCUSSION

### 4.1 Miscanthus and turfgrass support more similar microbiomes than maize

In this study, we explored the relationship between soil microbial communities, soil aggregation, and plant cover. We have previously shown at these same sites that miscanthus and turfgrass soils exhibit greater aggregate stability and maximum water-holding capacity than maize (29, 30). Here, we characterized bacterial and AMF communities across these three plant covers to ask whether microbiome composition co-varied with the observed structural differences and if so, whether plant cover identity or perennial life history better explained the patterns. Beta diversity across miscanthus, turfgrass, and maize revealed both a shared separation of the perennial covers from maize and a distinct plant-specific signal within the perennial treatments, highlighting the contributions of both life history and host identity to community structure. This observation aligns with previous studies that have linked key aspects of perennial cropping systems—permanent soil cover, no-till management, and increased root-derived carbon sources—to measurable shifts in microbial community composition (49, 50). Both bacterial and fungal communities are jointly shaped by plant selection and management, with roots identified as a significant moderating influence under perennial crops (51–53). These compositional differences between annual and perennial microbiomes may both reflect and reinforce functional differences, especially in relation to soil structural properties.

We found that aggregate stability and water-holding capacity differed among miscanthus, turfgrass, and maize, and that variation in soil structural properties aggregate stability, bulk density, and maximum water-holding capacity explained differences in microbiome structure across all three plant covers. This result is supported by previous observations that soil bacterial and fungal communities within microaggregates are shaped by long-term management activity in perennial tallgrass prairie (54, 55). Notably, while both miscanthus and turfgrass exhibited greater aggregate stability than maize, their microbiomes were not identical, pointing to plant cover identity, not perenniality alone, as a mediator of the relationship between soil structure and community composition. Distinct microbiomes tied to soil structural heterogeneity may also reflect functional differentiation: microbial communities in soils with greater aggregation have been associated with increased nutrient cycling, carbon stabilization, and resistance to drought stress (56, 57).

Given the role of plant cover identity in shaping the microbiome structure, we also expected microbial networks to differ not only between maize and the perennial crops but also miscanthus and turfgrass. To test this, we analyzed co-occurrence networks encompassing bacteria-bacteria, bacteria-AMF, and AMF-AMF interactions for taxa significantly correlated with water-stable aggregates. Overall, we found that the bacterial and fungal networks of the perennial plant covers in this study were significantly more interconnected than those in the annual maize system. Previous studies comparing microbial network complexity under annual and perennial plant covers generally find a similar pattern of enhanced inter-connectivity in perennials (58, 59), though other studies have reported contrasting findings (60), underscoring the need to consider management practices and site-specific context across diverse research settings. Heightened complexity in perennial-associated microbial communities may result from deep, branching root systems with continuous living roots, in contrast to the short-lived root architecture of annuals (3, 61). These longer interaction timeframes in perennials may promote sustained resource partitioning among cooccurring taxa, as organisms have more time to develop complementary ecological strategies (48, 62). While these mechanisms may generally explain increased complexity in perennials, plant-specific traits appear to shape the structure of microbial networks across different perennial plant covers.

Comparing the two perennial plant covers in our study, bacteria-bacteria interactions were most enriched in miscanthus, while AMF-AMF networks were most enriched in turfgrass (Figure 5). Although the two perennial treatments in this study had similar effects on aggregation, they differed significantly in soil biochemical measurements. Specifically, miscanthus soils were enriched in predominantly plant-derived carbohydrate monomers (arabinose and xylose). In contrast, turfgrass soils were enriched in predominantly microbial-associated monomers galactose and mannose (63). An open question is whether these biochemical differences reflect divergent resource-acquisition pathways in bacteria and AMF, manifesting in unique patterns of microbial community assembly tied to soil structural outcomes.

The inconsistent observations of fungal network patterns highlight the fact that AMF remain under-characterized in miscanthus and support the rationale for studying AMF in perennial systems. While this study represents three sites across a broad range of soil properties, we acknowledge the limitation of a single time point. It is possible that this single sampling time does not accurately capture temporal dynamics or the variable environmental contexts that may be important for studying AMF and its interactions. Given AMF’s known role in aggregation, targeted studies across multiple perennial systems are needed to clarify the functional roles of fungi and validate network-inferred interactions. Regardless, our results demonstrate that interaction network structure differs across miscanthus, turfgrass, and maize in ways that likely contribute to divergent outcomes in soil aggregation and underscore the need for targeted studies to clarify the mechanisms driving these plant-cover-specific patterns.

### 4.2 An enriched guild of “microbial architects” may promote aggregate stabilization under perennial plants (particularly miscanthus)

We narrowed our focus to a subset of microbes that could plausibly be associated with soil structural improvements. These putative “microbial architects” were identified based on whether the abundance of a given taxon was at least moderately correlated with soil aggregate stability. Of the 61 bacterial ASVs identified, many have been previously studied in the context of improvements in soil aggregation (Table 5). Specifically, these ASVs have been linked to aggregation (64), EPS production (65), and agricultural practices that promote soil health (66). Among the most enriched ASVs is one most similar to the genus *Actinophytocola,* which was previously found to be enriched in alfalfa soils and to show increased aggregate stability following bio-organic fertilizer amendment (67). Similarly, a dominant ASV most similar to genus *Pseudolabrys* was previously linked to soil aggregation driven by an organic biostimulant in citrus fields (68). Additionally, many of the observed aggregate-associated ASVs in this study are most similar to genera connected with well-studied plant-growth-promoting rhizobacteria. For example, the bacterial genus *Pseudomonas* is known for its prolific EPS production (5, 69) and *Nitrospira* regulates nitrification (70, 71).

**Table 5.**
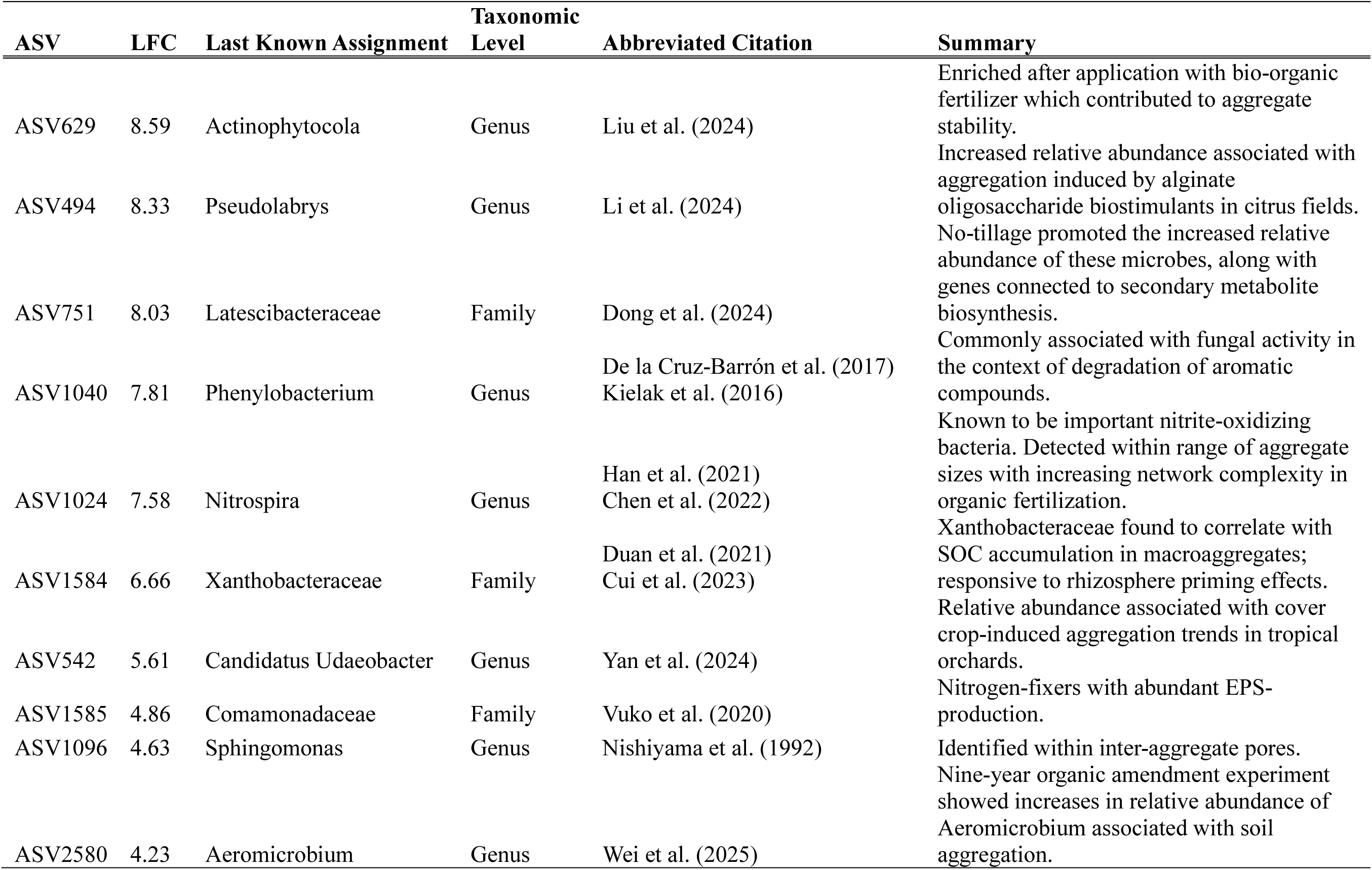
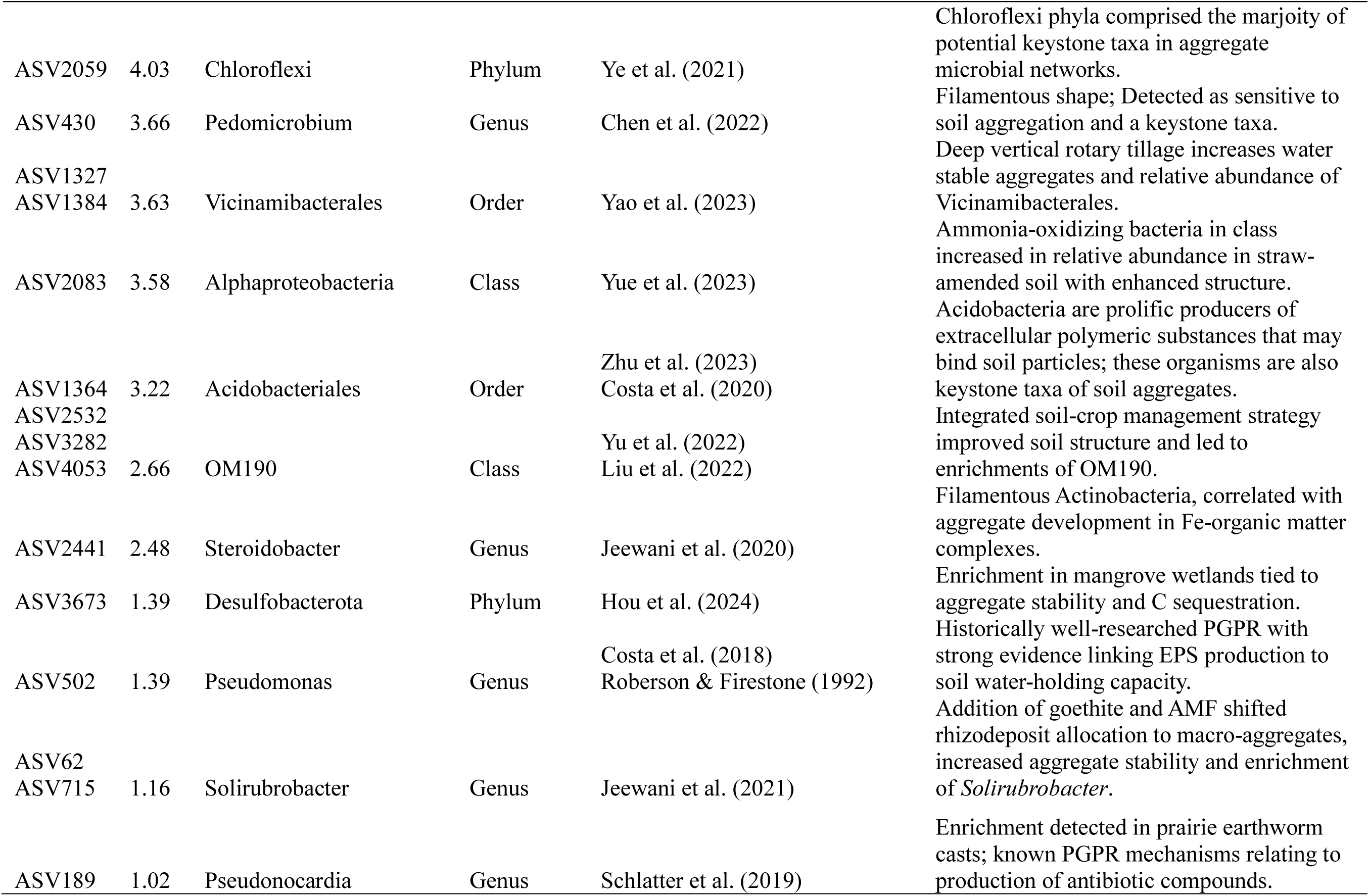
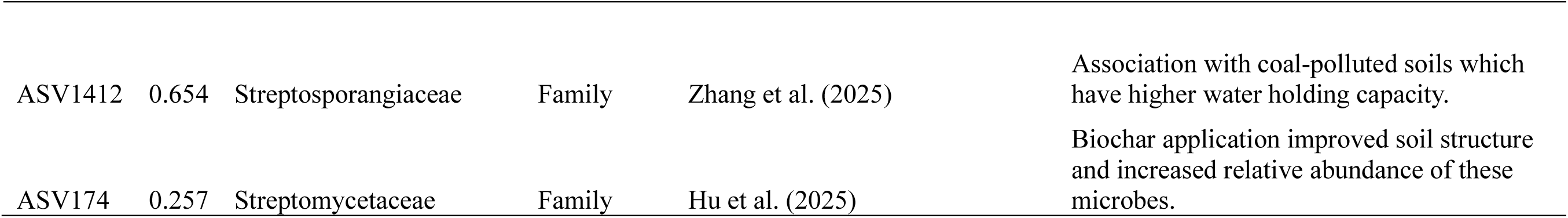
Summary of relevant literature on aggregation-associated taxa, listed by descending order of log2-fold change (LFC) in perennial vs. annual systems. Based on a Web of Science database query using the keywords (“soil aggregat*” AND “[name of taxa]”). Full citations can be found in the Supplemental Materials.

Fungi-bacteria interactions have also been identified as possible contributors to soil aggregation, and we detected evidence for potential relationships among our aggregate-correlated taxa. For example, bacteria in the genus *Solirubrobacter* were detected in hyphal-aggregate mineral associations with AMF (72) and *Phenylobacterium* is commonly associated with co-metabolism of aromatic compounds alongside fungi (73, 74). Generally, soil microorganisms rely on complex networks for nutrient acquisition, biofilm formation for protection against desiccation, and defense against antagonists, and bacterial-fungal interactions are likely involved in soil aggregation benefits.

If a guild of perennial-associated architect taxa were involved in soil aggregation, we would expect to find them consistently across perennial systems regardless of geographic location or sampling time. Thus, we cross-validated the presence of these putative architect taxa with three related publicly available datasets of miscanthus soil microbiomes across two sites (Urbana, Illinois, and Boone, Iowa) and two timepoints (2018 and 2022). We find that the putative bacterial architect taxa were highly prevalent across sites and over time (Figure 2). These results align with previous studies that have characterized stable miscanthus-associated bacterial communities over multiple site-years (23). In contrast, putative AMF architect taxa were fewer in number and less consistently observed across studies. This observation is likely influenced by the challenges and biases of AMF amplicon profiling (75), as well as discrete natural ranges and high variability in mycorrhizal fungal communities (76). However, half of the aggregate-correlated ASVs in our study were detected in the Illinois comparison site, which supports previous AMF-focused studies in miscanthus that have characterized communities of abundant, persistent, and plant-specific AMF taxa (77). For both putative bacteria and AMF architect taxa, their broad spatial and temporal distributions suggest they may be promising targets for further study regarding their links to aggregate stability.

Our approach identified putative architect taxa as possible microbial drivers of soil aggregation dynamics; however, these results are limited to observed correlations and require further mechanistic and functional studies to demonstrate direct linkages between microbial activity and aggregate formation. Further, it remains to be determined whether these bacteria or AMF can actively stabilize soil aggregates without direct plant involvement. An alternative hypothesis is that these taxa may merely be well-adapted to living in dense, root-structured environments that perennials provide. We suggest that these microorganisms exist along a spectrum of reciprocal plant-microbe interactions, where plant-beneficial activities of these taxa (e.g., nitrogen mineralization, EPS production, pathogen suppression) are stimulated by signaling from specific root exudates, or more broadly, rhizodeposited carbon (78, 79). Future experiments could be designed to elucidate these tradeoffs. For example, controlled incubations or greenhouse experiments could clarify the relative contributions of plant and microbial activity to aggregate stability. Additionally, metabolomic profiling of root exudates could help link rhizodeposition to differential microbial recruitment and aggregate formation.

### 4.3 Conclusion

Management of soil-plant-microbe interactions has the potential to help biomass crops sustain high productivity while also restoring degraded soils. *Miscanthus* × *giganteus* achieves this balance in part through improved soil aggregation and water-holding capacity, outcomes we show are linked to specific bacterial and AMF communities and putative architect taxa. Our study across three sites revealed that soil structural properties and microbial composition co-vary with plant cover identity across miscanthus, turfgrass, and maize, and that enriched specific bacterial and AMF taxa are positively correlated with aggregate stability. In miscanthus, we observe that bacteria-bacteria interactions among taxa related to soil aggregation are enhanced relative to those in maize and even in the perennial turfgrass. While more research is needed to understand the extent to which soil microbial communities facilitate soil aggregation and, in turn, water retention and subsequent biomass growth, our findings suggest that farmers can leverage soil-plant-microbe interactions to restore healthy soils and produce economically viable yields.

## FUNDING

This work was funded by the DOE Center for Advanced Bioenergy and Bioproducts Innovation (US Department of Energy, Office of Science, Biological and Environmental Research Program under Award Number DE-SC0018420). Any opinions, findings, and conclusions or recommendations expressed in this publication are those of the author(s) and do not necessarily reflect the views of the US Department of Energy.

## COMPETING INTERESTS

The authors have no relevant financial or non-financial interests to declare.

## AUTHOR CONTRIBUTIONS

**Phillip de Lorimier**: Formal analysis; investigation; methodology; software; validation; visualization; writing–original draft; writing–review and editing. **Jessica T. Nelson**: Investigation; methodology; writing–review and editing. **Bolívar Aponte Rolón**: Methodology; software; writing–review and editing. **Jared Flater**: Investigation; methodology; writing–review and editing. **Lorien Radmer**: Investigation; methodology; writing–review and editing. **Marshall D. McDaniel**: Conceptualization; funding acquisition; methodology; project administration; resources; supervision; visualization; writing–review and editing. **Adina Howe**: Conceptualization; funding acquisition; methodology; project administration; resources; supervision; visualization; writing–review and editing.

## ACKNOWLEDGEMENTS

We thank Nicholas Boersma, Andy VanLoocke, and Emily Heaton for initiating and maintaining the LAMPS field sites. We thank Mareli Sanchez Julia for assistance with AMF amplicon analysis and Benjamin Jacobs for statistical analysis consulting. We thank the members of the McDaniel and Howe labs for constructive comments during the preparation of the manuscript.

## Notes

### Competing Interest Statement

The authors have declared no competing interest.

