## Supplemental Materials for "Could microbes be the architects of improved soil structure under *Miscanthus* × *giganteus*?"

for

TABLE OF CONTENTS

SUPPLEMENTAL TABLES

1. **Table S1.** Description of soil physical measurements referenced in this manuscript	2
2. **Table S2.** Taxonomic classification of bacterial and arbuscular mycorrhizal fungal amplicon sequence variants (ASVs) positively correlated with soil aggregate stability	3
3. **Table S3.** Network analysis metrics describing within-module connectivity (zi) and between-module connectivity (pi)	6
4. **Table S4.** Full citations for references cited in Table 5	7

SUPPLEMENTAL FIGURES

1. **Figure S1.** Water-stable aggregate measurement of soil samples in this study	9
2. **Figure S2.** Illustration of potential direct and indirect pathways contributing to the observed increase in soil maximum water holding capacity under *Miscanthus* × *giganteus*	10
3. **Figure S3.** Counts of unique and shared amplicon sequence variants	11
4. **Figure S4.** Taxonomic distribution of the most abundant microbial taxa at the level of genus	12
5. **Figure S5.** Measurements of indicator genes for microbial groups by quantitative PCR	13

**Table S1**. Soil physical measurements used in this manuscript and summarized here. See Nelson et al. (2026) for more details.

| Soil Measurement | Abbreviation | Units | Brief Methodological Description | Corresponding Reference |
| --- | --- | --- | --- | --- |
| Bulk Density | BD | g cm^-3^ | Calculated from sample mass and volume dimensions of HYPROP sample ring. | Singh & Verdi (2024) |
| Root Density | RD | g cm^-3^ | Density of roots in the top 5 cm of soil. |  |
| Soil Organic Matter | SOM | % | Loss on ignition (LOI) method. |  |
| Water Stable Aggregates | WSA | % | Wet-sieve aggregates through nested sieves and calculate proportion of soil mass remaining in sieves >0.25 mm (“stable aggregates”). | Obrycki et al. (2018) |
| Penetration Resistance |  | kPa | Manual push penetrometer. |  |
| Soil water retention |  | % | Determined water content at varying matric potentials from HYPROP evaporation method using tensiometer and balance measurements. | Singh & Verdi (2024) |
| Maximum water holding capacity | MWHC | % | Using the funnel-filter paper method and drained for 6 h | Nelson et al. (2024) |
| Microbial Biomass Carbon | MBC | mg kg^-1^ | Chloroform fumigation-extraction to lyse microbial cells, extracted with K₂SO₄. | Joergensen (1996) |
| Salt-Extractable Organic Carbon | SEOC | mg kg^-1^ | Non-fumigated samples were extracted in parallel using K₂SO₄. | Joergensen (1996) |
| Extracellular Polymeric Substances | EPS | mg kg^-1^ | Extracted using cation exchange resin method. | Redmile-Gordon et al. (2014) |
| Soil Carbohydrates |  | mg kg^-1^ | Sequential acid hydrolysis and quantified with ion exchange chromatography with pulsed amperometric detection. | Olk (2008) |

**Table S2**. Taxonomic classification of bacterial and arbuscular mycorrhizal fungal amplicon sequence variants (ASVs) positively correlated with soil aggregate stability (𝜌).

| ASV | 𝜌 | Kingdom | Phylum | Class | Order | Family | Genus | Species |
| --- | --- | --- | --- | --- | --- | --- | --- | --- |
| ASV32 | 0.463 | Bacteria | Actinobacteriota | Actinobacteria | Micrococcales | Intrasporangiaceae | Knoellia | - |
| ASV33 | 0.424 | Bacteria | Actinobacteriota | Thermoleophilia | Solirubrobacterales | Solirubrobacteraceae | Conexibacter | - |
| ASV46 | 0.437 | Bacteria | Actinobacteriota | Actinobacteria | Frankiales | Frankiaceae | Jatrophihabitans | - |
| ASV55 | 0.541 | Bacteria | Proteobacteria | Gammaproteobacteria | Xanthomonadales | Rhodanobacteraceae | - | - |
| ASV57 | 0.497 | Bacteria | Proteobacteria | Gammaproteobacteria | Burkholderiales | TRA3-20 | - | - |
| ASV62 | 0.425 | Bacteria | Actinobacteriota | Thermoleophilia | Solirubrobacterales | Solirubrobacteraceae | Solirubrobacter | - |
| ASV63 | 0.408 | Bacteria | Bacteroidota | Bacteroidia | Chitinophagales | Chitinophagaceae | Puia | - |
| ASV72 | 0.466 | Bacteria | Actinobacteriota | Thermoleophilia | Solirubrobacterales | 67-14 | - | - |
| ASV99 | 0.433 | Bacteria | Chloroflexi | Ktedonobacteria | Ktedonobacterales | JG30-KF-AS9 | - | - |
| ASV137 | 0.43 | Bacteria | Proteobacteria | Alphaproteobacteria | Micropepsales | Micropepsaceae | - | - |
| ASV150 | 0.404 | Bacteria | Acidobacteriota | Acidobacteriae | Bryobacterales | Bryobacteraceae | Bryobacter | - |
| ASV155 | 0.415 | Bacteria | Verrucomicrobiota | Verrucomicrobiae | Pedosphaerales | Pedosphaeraceae | ADurb.Bin063-1 | - |
| ASV174 | 0.431 | Bacteria | Actinobacteriota | Actinobacteria | Streptomycetales | Streptomycetaceae | - | - |
| ASV189 | 0.411 | Bacteria | Actinobacteriota | Actinobacteria | Pseudonocardiales | Pseudonocardiaceae | Pseudonocardia | - |
| ASV229 | 0.423 | Bacteria | Proteobacteria | Gammaproteobacteria | Burkholderiales | SC-I-84 | - | - |
| ASV237 | 0.458 | Bacteria | Acidobacteriota | Subgroup 5 | - | - | - | - |
| ASV265 | 0.486 | Bacteria | Acidobacteriota | Acidobacteriae | Solibacterales | Solibacteraceae | Candidatus Solibacter | - |
| ASV267 | 0.438 | Bacteria | Actinobacteriota | Actinobacteria | Frankiales | Acidothermaceae | Acidothermus | - |
| ASV271 | 0.436 | Bacteria | Proteobacteria | Gammaproteobacteria | Xanthomonadales | Rhodanobacteraceae | Dokdonella | - |
| ASV391 | 0.506 | Bacteria | Proteobacteria | Gammaproteobacteria | CCD24 | - | - | - |
| ASV396 | 0.498 | Bacteria | Myxococcota | Polyangia | Polyangiales | BIrii41 | - | - |
| ASV406 | 0.407 | Bacteria | Myxococcota | Polyangia | Polyangiales | BIrii41 | - | - |
| ASV430 | 0.456 | Bacteria | Proteobacteria | Alphaproteobacteria | Rhizobiales | Hyphomicrobiaceae | Pedomicrobium | - |
| ASV463 | 0.45 | Bacteria | Proteobacteria | Alphaproteobacteria | Rhizobiales | KF-JG30-B3 | - | - |
| ASV486 | 0.421 | Bacteria | Actinobacteriota | Actinobacteria | Frankiales | Acidothermaceae | Acidothermus | - |
| ASV494 | 0.417 | Bacteria | Proteobacteria | Alphaproteobacteria | Rhizobiales | Xanthobacteraceae | Pseudolabrys | - |
| ASV502 | 0.46 | Bacteria | Proteobacteria | Gammaproteobacteria | Pseudomonadales | Pseudomonadaceae | Pseudomonas | - |
| ASV542 | 0.402 | Bacteria | Verrucomicrobiota | Verrucomicrobiae | Chthoniobacterales | Chthoniobacteraceae | Candidatus Udaeobacter | - |
| ASV560 | 0.409 | Bacteria | Chloroflexi | Ktedonobacteria | Ktedonobacterales | Ktedonobacteraceae | - | - |
| ASV566 | 0.458 | Bacteria | Myxococcota | Polyangia | Polyangiales | BIrii41 | - | - |
| ASV629 | 0.418 | Bacteria | Actinobacteriota | Actinobacteria | Pseudonocardiales | Pseudonocardiaceae | Actinophytocola | - |
| ASV640 | 0.467 | Bacteria | Bacteroidota | Bacteroidia | Chitinophagales | Chitinophagaceae | Parafilimonas | - |
| ASV665 | 0.467 | Bacteria | Actinobacteriota | Thermoleophilia | Gaiellales | - | - | - |
| ASV715 | 0.488 | Bacteria | Actinobacteriota | Thermoleophilia | Solirubrobacterales | Solirubrobacteraceae | Solirubrobacter | - |
| ASV751 | 0.403 | Bacteria | Latescibacterota | Latescibacteria | Latescibacterales | Latescibacteraceae | - | - |
| ASV777 | 0.452 | Bacteria | Proteobacteria | Gammaproteobacteria | Xanthomonadales | Rhodanobacteraceae | Rhodanobacter | - |
| ASV827 | 0.411 | Bacteria | Actinobacteriota | Thermoleophilia | Gaiellales | - | - | - |
| ASV905 | 0.436 | Bacteria | Acidobacteriota | Acidobacteriae | Acidobacteriales | Koribacteraceae | Candidatus Koribacter | - |
| ASV1009 | 0.411 | Bacteria | Actinobacteriota | Actinobacteria | Micromonosporales | Micromonosporaceae | - | - |
| ASV1024 | 0.43 | Bacteria | Nitrospirota | Nitrospiria | Nitrospirales | Nitrospiraceae | Nitrospira | - |
| ASV1040 | 0.423 | Bacteria | Proteobacteria | Alphaproteobacteria | Caulobacterales | Caulobacteraceae | Phenylobacterium | - |
| ASV1078 | 0.462 | Bacteria | Acidobacteriota | Holophagae | Subgroup 7 | - | - | - |
| ASV1096 | 0.415 | Bacteria | Proteobacteria | Alphaproteobacteria | Sphingomonadales | Sphingomonadaceae | Sphingomonas | - |
| ASV1111 | 0.537 | Bacteria | Actinobacteriota | Thermoleophilia | - | - | - | - |
| ASV1139 | 0.519 | Bacteria | Actinobacteriota | Actinobacteria | Frankiales | Acidothermaceae | Acidothermus | - |
| ASV1175 | 0.415 | Bacteria | Verrucomicrobiota | Verrucomicrobiae | Chthoniobacterales | Chthoniobacteraceae | Chthoniobacter | - |
| ASV1185 | 0.432 | Bacteria | Actinobacteriota | Thermoleophilia | Solirubrobacterales | 67-14 | - | - |
| ASV1323 | 0.44 | Bacteria | Proteobacteria | Alphaproteobacteria | Acetobacterales | Acetobacteraceae | - | - |
| ASV1351 | 0.498 | Bacteria | Actinobacteriota | Acidimicrobiia | IMCC26256 | - | - | - |
| ASV1384 | 0.417 | Bacteria | Acidobacteriota | Vicinamibacteria | Vicinamibacterales | - | - | - |
| ASV1479 | 0.407 | Bacteria | Actinobacteriota | Acidimicrobiia | Microtrichales | Ilumatobacteraceae | CL500-29 marine group | - |
| ASV1537 | 0.481 | Bacteria | Verrucomicrobiota | Verrucomicrobiae | Pedosphaerales | Pedosphaeraceae | - | - |
| ASV1584 | 0.501 | Bacteria | Proteobacteria | Alphaproteobacteria | Rhizobiales | Xanthobacteraceae | - | - |
| ASV2059 | 0.452 | Bacteria | Chloroflexi | - | - | - | - | - |
| ASV2083 | 0.408 | Bacteria | Proteobacteria | Alphaproteobacteria | - | - | - | - |
| ASV2471 | 0.409 | Bacteria | Myxococcota | Polyangia | Nannocystales | Nannocystaceae | Nannocystis | - |
| ASV2843 | 0.449 | Bacteria | Armatimonadota | - | - | - | - | - |
| ASV3615 | 0.44 | Bacteria | Proteobacteria | Alphaproteobacteria | Rickettsiales | Mitochondria | - | - |
| ASV3673 | 0.48 | Bacteria | Desulfobacterota | - | - | - | - | - |
| ASV3804 | 0.436 | Bacteria | Proteobacteria | Alphaproteobacteria | Acetobacterales | Acetobacteraceae | Rhodovastum | - |
| ASV4726 | 0.413 | Bacteria | Proteobacteria | Alphaproteobacteria | Rickettsiales | Mitochondria | - | - |
| ASV6 | 0.412 | Fungi | Mucoromycota | Glomeromycetes | Glomerales | Glomeraceae | Glomus | VTX00064 |
| ASV95 | 0.433 | Fungi | Mucoromycota | Glomeromycetes | Glomerales | Glomeraceae | Glomus | VTX00151 |
| ASV95 | 0.417 | Fungi | Mucoromycota | Glomeromycetes | Glomerales | Glomeraceae | Glomus | VTX00151 |
| ASV131 | 0.459 | Fungi | Mucoromycota | Glomeromycetes | Glomerales | Glomeraceae | Glomus | VTX00212 |
| ASV131 | 0.436 | Fungi | Mucoromycota | Glomeromycetes | Glomerales | Glomeraceae | Glomus | VTX00212 |
| ASV132 | 0.443 | Fungi | Mucoromycota | Glomeromycetes | Glomerales | Glomeraceae | Glomus | VTX00423 |
| ASV194 | 0.432 | Fungi | Mucoromycota | Glomeromycetes | Glomerales | Glomeraceae | Glomus | VTX00212 |
| ASV194 | 0.416 | Fungi | Mucoromycota | Glomeromycetes | Glomerales | Glomeraceae | Glomus | VTX00212 |
| ASV205 | 0.432 | Fungi | Mucoromycota | Glomeromycetes | Glomerales | Glomeraceae | Glomus | VTX00151 |
| ASV205 | 0.416 | Fungi | Mucoromycota | Glomeromycetes | Glomerales | Glomeraceae | Glomus | VTX00151 |
| ASV329 | 0.416 | Fungi | Mucoromycota | Glomeromycetes | Glomerales | Glomeraceae | Glomus | - |
| ASV395 | 0.402 | Fungi | Mucoromycota | Glomeromycetes | Glomerales | - | - | - |

Table S3. Network analysis metrics describing within-module connectivity (zi) and between-module connectivity (pi). Pairwise treatment comparisons for each response variable and network type, restricted to nodes correlated with water-stable aggregates (WSA). For Bacteria-AMF and AMF-AMF networks, only 16S ASVs were filtered for WSA correlation; AMF nodes were retained regardless of WSA correlation status. Estimates and standard errors are from linear models with Treatment as a fixed effect. P-values adjusted using Tukey method.

| **Metric** | **Network** | **Contrast** | **Estimate** | **Standard error** | **Degrees of freedom** | **t-ratio** | **p-value** |
| --- | --- | --- | --- | --- | --- | --- | --- |
| Pi | Bacteria-Bacteria | Maize - Mxg | -0.048 | 0.029 | 280 | -1.653 | 0.225 |
|  |  | Maize - Turf | -0.032 | 0.030 | 280 | -1.063 | 0.538 |
|  |  | Mxg - Turf | 0.016 | 0.025 | 280 | 0.658 | 0.788 |
|  | Bacteria-AMF | Maize - Mxg | 0.030 | 0.021 | 496 | 1.423 | 0.330 |
|  |  | Maize - Turf | 0.012 | 0.020 | 496 | 0.582 | 0.830 |
|  |  | Mxg - Turf | -0.018 | 0.021 | 496 | -0.859 | 0.667 |
|  | AMF-AMF | Maize - Mxg | 0.032 | 0.018 | 252 | 1.778 | 0.179 |
|  |  | Maize - Turf | -0.007 | 0.015 | 252 | -0.483 | 0.879 |
|  |  | Mxg - Turf | -0.039 | 0.019 | 252 | -2.079 | 0.096 |
| Zi | Bacteria-Bacteria | Maize - Mxg | 0.106 | 0.144 | 280 | 0.736 | 0.743 |
|  |  | Maize - Turf | 0.116 | 0.148 | 280 | 0.786 | 0.712 |
|  |  | Mxg - Turf | 0.010 | 0.123 | 280 | 0.083 | 0.996 |
|  | Bacteria-AMF | Maize - Mxg | 0.601 | 0.293 | 496 | 2.052 | 0.101 |
|  |  | Maize - Turf | 0.640 | 0.280 | 496 | 2.283 | 0.059 |
|  |  | Mxg - Turf | 0.040 | 0.295 | 496 | 0.135 | 0.990 |
|  | AMF-AMF | Maize - Mxg | 0.000 | 0.190 | 195 | 0.000 | 1.000 |
|  |  | Maize - Turf | 0.000 | 0.156 | 195 | 0.000 | 1.000 |
|  |  | Mxg - Turf | 0.000 | 0.193 | 195 | 0.000 | 1.000 |

**Table S4.** Full citations for references cited in Table 5.

| **Abbreviated Citation** | **Full Citation** |
| --- | --- |
| Chen et al. (2022) | Chen J, Song D, Liu D et al. Soil Aggregation Shaped the Distribution and Interaction of Bacterial-Fungal Community Based on a 38-Year Fertilization Experiment in China. Front Microbiol 2022;13. https://doi.org/10.3389/fmicb.2022.824681. |
| Costa et al. (2018) | Costa OYA, Raaijmakers JM, Kuramae EE. Microbial Extracellular Polymeric Substances: Ecological Function and Impact on Soil Aggregation. Front Microbiol 2018;9. https://doi.org/10.3389/fmicb.2018.01636. |
| Costa et al. (2020) | Costa OYA, Pijl A, Kuramae EE. Dynamics of active potential bacterial and fungal interactions in the assimilation of acidobacterial EPS in soil. Soil Biology and Biochemistry 2020;148:107916. https://doi.org/10.1016/j.soilbio.2020.107916. |
| Cui et al. (2023) | Cui H, Chen P, He C et al. Soil microbial community structure dynamics shape the rhizosphere priming effect patterns in the paddy soil. Science of The Total Environment 2023;857:159459. https://doi.org/10.1016/j.scitotenv.2022.159459. |
| De la Cruz-Barrón et al. (2017) | De la Cruz-Barrón M, Cruz-Mendoza A, Navarro–Noya YE et al. The Bacterial Community Structure and Dynamics of Carbon and Nitrogen when Maize (Zea mays L.) and Its Neutral Detergent Fibre Were Added to Soil from Zimbabwe with Contrasting Management Practices. Microb Ecol 2017;73(1):135–52. https://doi.org/10.1007/s00248-016-0807-8. |
| Dong et al. (2024) | Dong F, Wang L, Xu T et al. Multi-omics analysis of soil microbiota and metabolites in dryland wheat fields under different tillage methods. Sci Rep 2024;14(1):24066. https://doi.org/10.1038/s41598-024-74620-0. |
| Duan et al. (2021) | Duan Y, Chen L, Li Y et al. N, P and straw return influence the accrual of organic carbon fractions and microbial traits in a Mollisol. Geoderma 2021;403:115373. https://doi.org/10.1016/j.geoderma.2021.115373. |
| Han et al. (2021) | Han S, Huang Q, Chen W. Partitioning Nitrospira community structure and co-occurrence patterns in a long-term inorganic and organic fertilization soil. J Soils Sediments 2021;21(2):1099–108. https://doi.org/10.1007/s11368-020-02813-x. |
| Hou et al. (2024) | Hou N, Yang X, Wang W et al. Mangrove wetland recovery enhances soil carbon sequestration capacity of soil aggregates and microbial network stability in southeastern China. Science of The Total Environment 2024;951:175586. https://doi.org/10.1016/j.scitotenv.2024.175586. |
| Hu et al. (2025) | Hu J, Yang S, Cornelis WM et al. Microstructure and Microorganisms Alternation of Paddy Soil: Interplay of Biochar and Water-Saving Irrigation. Plants 2025;14(10):1498. https://doi.org/10.3390/plants14101498. |
| Jeewani et al. (2020) | Jeewani PH, Gunina A, Tao L et al. Rusty sink of rhizodeposits and associated keystone microbiomes. Soil Biology and Biochemistry 2020;147:107840. https://doi.org/10.1016/j.soilbio.2020.107840. |
| Jeewani et al. (2021) | Jeewani PH, Luo Y, Yu G et al. Arbuscular mycorrhizal fungi and goethite promote carbon sequestration via hyphal-aggregate mineral interactions. Soil Biology and Biochemistry 2021;162:108417. https://doi.org/10.1016/j.soilbio.2021.108417. |
| Kielak et al. (2016) | Kielak AM, Scheublin TR, Mendes LW et al. Bacterial Community Succession in Pine-Wood Decomposition. Front Microbiol 2016;7. https://doi.org/10.3389/fmicb.2016.00231. |
| Li et al. (2024) | Li Z, Duan S, Ouyang X et al. Coupled soil moisture management and alginate oligosaccharide strategies enhance citrus orchard production, water and potassium use efficiency by improving the rhizosphere soil environment. Agricultural Water Management 2024;297:108828. https://doi.org/10.1016/j.agwat.2024.108828. |
| Liu et al. (2022) | Liu X, Liu H, Ren D et al. Interlinkages between soil properties and keystone taxa under different tillage practices on the North China Plain. Applied Soil Ecology 2022;178:104551. https://doi.org/10.1016/j.apsoil.2022.104551. |
| Liu et al. (2024) | Liu T, Wang Q, Li Y et al. Bio-organic fertilizer facilitated phytoremediation of heavy metal(loid)s-contaminated saline soil by mediating the plant-soil-rhizomicrobiota interactions. Science of The Total Environment 2024;922:171278. https://doi.org/10.1016/j.scitotenv.2024.171278. |
| Nishiyama et al. (1992) | Nishiyama M, Senoo K, Wada H et al. Identification of soil micro-habitats for growth, death and survival of a bacterium, γ-1,2,3,4,5,6-hexachlorocyclohexane-assimilating Sphingomonas paucimobilis, by fractionation of soil. FEMS Microbiol Ecol 1992;10(3):145–50. https://doi.org/10.1111/j.1574-6941.1992.tb01650.x. |
| Roberson & Firestone (1992) | Roberson EB, Firestone MK. Relationship between Desiccation and Exopolysaccharide Production in a Soil Pseudomonas sp. Applied and Environmental Microbiology 1992;58(4):1284–91. https://doi.org/10.1128/aem.58.4.1284-1291.1992. |
| Schlatter et al. (2019) | Schlatter DC, Baugher CM, Kahl K et al. Bacterial communities of soil and earthworm casts of native Palouse Prairie remnants and no-till wheat cropping systems. Soil Biology and Biochemistry 2019;139:107625. https://doi.org/10.1016/j.soilbio.2019.107625. |
| Vuko et al. (2020) | Vuko M, Cania B, Vogel C et al. Shifts in reclamation management strategies shape the role of exopolysaccharide and lipopolysaccharide-producing bacteria during soil formation. Microbial Biotechnology 2020;13(2):584–98. https://doi.org/10.1111/1751-7915.13532. |
| Wei et al. (2025) | Wei B, Bi J, Qian X et al. Organic Manure Amendment Fortifies Soil Health by Enriching Beneficial Metabolites and Microorganisms and Suppressing Plant Pathogens. Agronomy 2025;15(2):429. https://doi.org/10.3390/agronomy15020429. |
| Yan et al. (2024) | Yan C, An D, Zhao B et al. Intercropping herbage promoted the availability of soil phosphorus and improved the bacterial genus structure and the abundance of key bacterial taxa in the acidic soil of mango (Mangifera indica L.) orchards. Soil Use and Management 2024;40(2):e13046. https://doi.org/10.1111/sum.13046. |
| Yao et al. (2023) | Yao R, Gao Q, Liu Y et al. Deep vertical rotary tillage mitigates salinization hazards and shifts microbial community structure in salt-affected anthropogenic-alluvial soil. Soil and Tillage Research 2023;227:105627. https://doi.org/10.1016/j.still.2022.105627. |
| Ye et al. (2021) | Ye G, Banerjee S, He JZ et al. Manure application increases microbiome complexity in soil aggregate fractions: Results of an 18-year field experiment. Agriculture, Ecosystems & Environment 2021;307:107249. https://doi.org/10.1016/j.agee.2020.107249. |
| Yu et al. (2022) | Yu N, Liu J, Ren B et al. Long-term integrated soil-crop management improves soil microbial community structure to reduce GHG emission and increase yield. Front Microbiol 2022;13. https://doi.org/10.3389/fmicb.2022.1024686. |
| Yue et al. (2023) | Yue Y, Gong X, Zheng Y et al. Organic Material Addition Optimizes Soil Structure by Enhancing Copiotrophic Bacterial Abundances of Nitrogen Cycling Microorganisms in Northeast China. Agronomy 2023;13(8):2108. https://doi.org/10.3390/agronomy13082108. |
| Zhang et al. (2025) | Zhang W, Nie X, Zhao T et al. The Impact of Coal Pollution on Soil Microbial Diversity and Community Structure. J Soil Sci Plant Nutr 2025;25(1):1706–22. https://doi.org/10.1007/s42729-025-02232-2. |
| Zhu et al. (2023) | Zhu K, Jia W, Mei Y et al. Shift from flooding to drying enhances the respiration of soil aggregates by changing microbial community composition and keystone taxa. Front Microbiol 2023;14. https://doi.org/10.3389/fmicb.2023.1167353. |

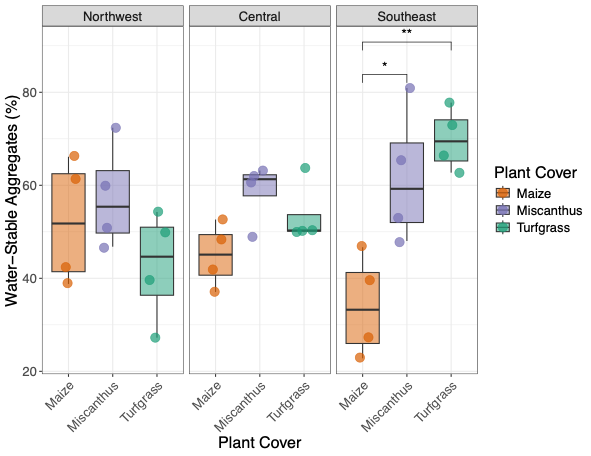

**Figure S1**. Water-stable aggregate percentage (proportion of soil mass remaining in sieves >0.25 mm) of soil samples comparing across plant covers and sites. Significant differences within site and among plant cover means are shown: * p ≤ 0.05; ** p ≤ 0.01; *** p ≤ 0.001. Adapted from Nelson et al. (2026).

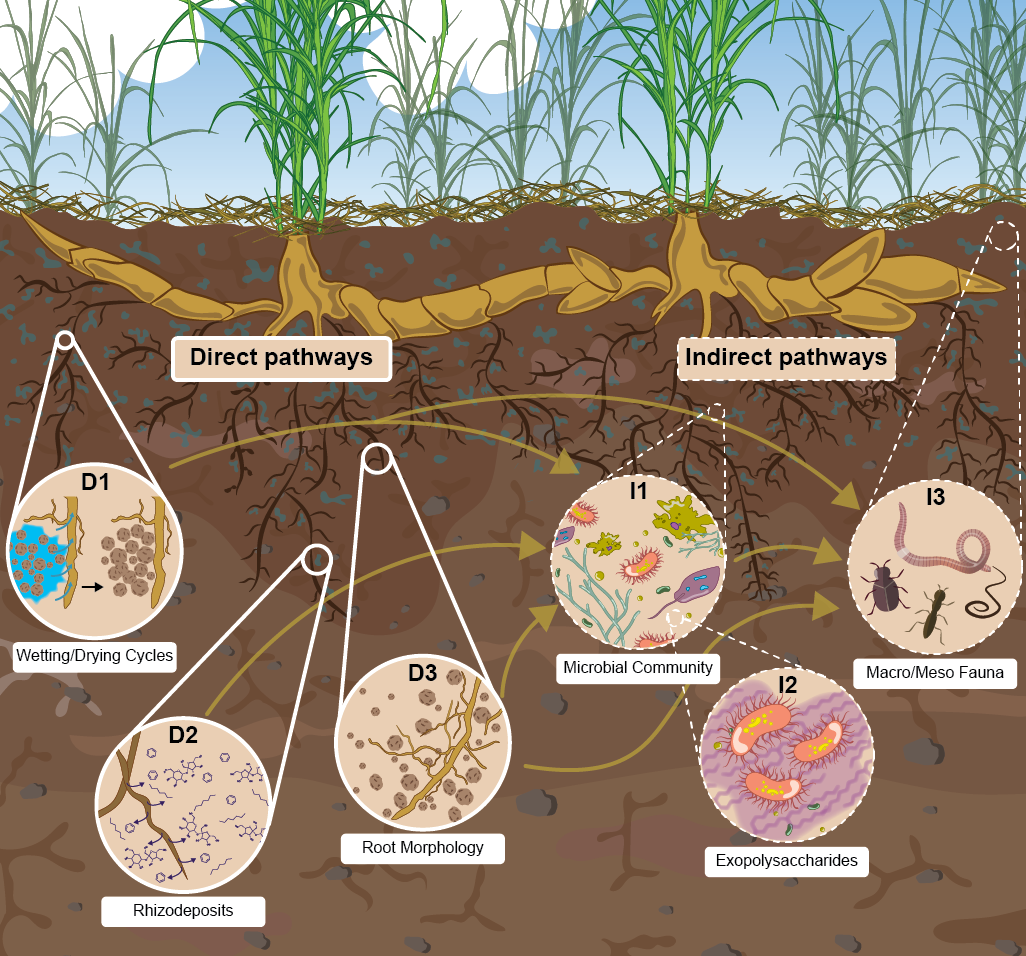

Figure S2. Illustration of potential direct and indirect pathways contributing to the observed increase in soil maximum water holding capacity under Miscanthus × giganteus. (Adapted from Nelson et al. (2025). Illustration by Kayla Nass).

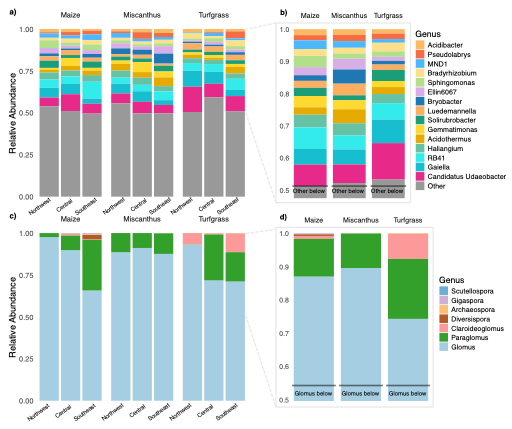

Figure S3. Taxonomic distribution of dominant microbial taxa at the level of genus. (a) Relative abundance of bacterial genera separated by plant cover and site. (b) Detail of most abundant bacterial genera aggregated by plant cover. (c) Relative abundance of arbuscular mycorrhizal fungal genera separated by plant cover and site. (d) Detail of most abundant arbuscular mycorrhizal fungal genera aggregated by plant cover.

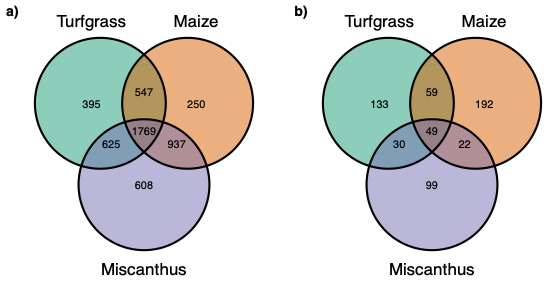

Figure S4. Counts of unique and shared amplicon sequence variants detected among treatments for (a) bacteria and (b) arbuscular mycorrhizal fungi.

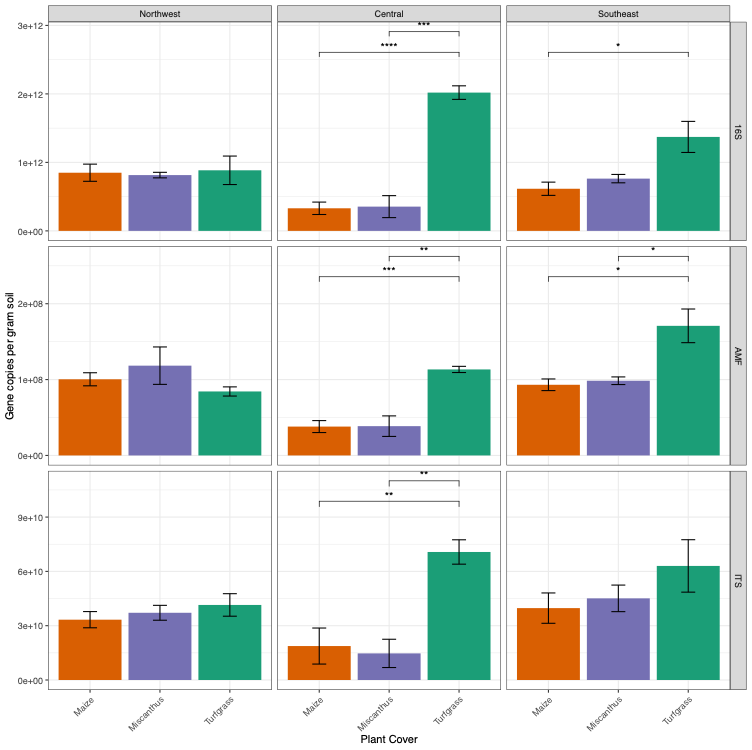

Figure S5. Measurements of indicator genes for microbial groups by quantitative PCR: (top) bacteria; (middle) arbuscular mycorrhizal fungi; (bottom) all fungi. Significant differences within site and among plant cover means are shown: * p ≤ 0.05; ** p ≤ 0.01; *** p ≤ 0.001.
